# A Semantic Framework for Predicting Herbal-Drug Biotransformation Conflicts via Biomedical Literature Mining

**DOI:** 10.1101/2025.08.07.668963

**Authors:** Kanchan Verandani

## Abstract

The widespread use of herbal supplements alongside conventional medicines increases the risk of unpredictable interactions affecting drug absorption and metabolism. This study introduces a comprehensive semantic framework that synthesizes knowledge from biomedical ontologies, curated databases, and full-text literature to model potential biotransformation conflicts between natural compounds and pharmaceutical agents. Leveraging advanced relation extraction systems and graph-based inference techniques, we constructed an enriched knowledge graph capable of highlighting mechanistic pathways involving enzymes, transporters, and drug constituents. Case studies with compounds like green tea and kratom demonstrate the framework’s potential to surface both known and previously underexplored interactions. The proposed system offers a scalable, hypothesis-generating platform for early-stage pharmacokinetic safety analysis in the context of natural product co-administration.

## I. Introduction

The utilization of botanical supplements and other naturally derived compounds has seen a marked increase following the enactment of the Dietary Supplement Health and Education Act of 1994. Approximately 18% of adults in the United States have reported consistent intake of such supplements, and sales of herbal-based dietary aids in the country rose by 9.7% in 2021. While these agents are not designed to substitute conventional medications, simultaneous consumption with prescription drugs is widespread, particularly among elderly populations, where concurrent usage reaches up to 88%.

This co-administration of pharmaceuticals and natural products may prompt pharmacokinetic interactions known as natural product-drug interactions (NPDIs), potentially altering drug efficacy or safety. NPDIs arise when natural constituents influence the pharmacokinetic parameters—absorption, distribution, metabolism, or excretion (ADME)—of medications, possibly attenuating therapeutic outcomes or eliciting adverse events.

For instance, green tea (*Camellia sinensis*), commonly consumed both as a beverage and a supplement, was observed to interfere with the antihypertensive drug nadolol. The proposed mechanism involves inhibition of intestinal organic anion transporting polypeptides (OATPs), which facilitate drug uptake into systemic circulation, thereby reducing nadolol’s efficacy. In contrast, some phytoconstituents may enhance drug absorption. Piperine, derived from black pepper (*Piper nigrum L*.), can elevate oral bioavailability of co-administered xenobiotics by inhibiting intestinal cytochrome P450 isoenzyme CYP3A4.

A comprehensive understanding of the molecular pathways that drive these interactions is essential for risk mitigation. Analogous to drug-drug interactions, computational methodologies can elucidate the mechanistic basis for NPDIs. Recent efforts have explored natural product-drug pairings using biomedical literature mining and construction of knowledge graphs (KGs) to predict plausible interactions.

Dietary supplements in the United States encompass a wide spectrum of botanical products regulated differently than synthetic pharmaceuticals. The U.S. Food and Drug Administration (FDA) lacks pre-market approval authority for these substances and may only act post-marketing in cases of demonstrated harm.

To forecast pharmacokinetic NPDIs, a workflow similar to drug interaction assessment is employed. Initially, *in vitro* experimentation determines whether specific natural compounds inhibit or induce drug-metabolizing enzymes or transporters. These outcomes are then translated into clinical predictions using in vitro-in vivo extrapolation (IVIVE), followed by physiologically based pharmacokinetic (PBPK) modeling or human trials to confirm relevance.

However, outcomes predicted through IVIVE do not always correlate with clinical findings. For example, although static modeling suggested green tea would increase systemic exposure to raloxifene via UDP-glucuronosyltransferase (UGT) inhibition, clinical studies revealed a contrary effect. Given green tea’s widespread consumption and claimed health benefits—ranging from cognitive enhancement to cardiovascular support—the risk of clinically significant NPDIs is non-trivial.

Consequently, devising intelligent frameworks for timely NPDI detection remains paramount. It was postulated that an ontologically grounded biomedical knowledge graph enriched with curated scientific literature could provide mechanistic insight into NPDIs. To validate this hypothesis, a dedicated knowledge graph, denoted as NP-KG, was developed. Its capacity to represent known pharmacokinetic mechanisms involving green tea and kratom (*Mitragyna speciosa*)—another widely used natural product—was examined. Kratom is frequently utilized for pain management, anxiety relief, and opioid withdrawal. The plant’s alkaloids possess psychoactive properties and raise concerns regarding toxicity and pharmacokinetic interference. NP-KG was further employed to derive mechanistic hypotheses for NPDIs related to kratom based on existing case studies.

## II. Materials and Methods

A biomedical knowledge graph (KG) encapsulates entities spanning diverse biological domains—such as genes, chemicals, disorders, and molecular processes—using nodes, while interrelations among these entities are represented via edges. This section outlines the ontology-based KG construction, semantic relation extraction from full-text scientific documents, and the assimilation of data into NP-KG. The relation extraction phase yields semantic triplets in the form `subject-relation-object`, where entities are modeled as nodes and the relationship becomes the connecting edge.

### A. Data Aggregation

Ontologies provide structured vocabularies representing specialized domain knowledge. These frameworks define classes—representing sets of real-world instances—and the relational properties connecting them. For example, the ChEBI (Chemical Entities of Biological Interest) ontology offers detailed definitions and hierarchical classifications for chemical substances, such as the flavonoid “catechin” identified as `ChEBI:23053`.

Biomedical ontologies encapsulate contemporary biological understanding and serve as a foundation for expressing knowledge statements within graph structures. The OBO Foundry maintains a suite of interoperable ontologies designed under shared guidelines, ensuring consistency, reusability, and logical structure.

To construct the NP-KG, the Phenotype Knowledge Translator (PheKnowLator, version 3.0.0) workflow was adopted. PheKnowLator is a Python-based framework for assembling large-scale semantically integrated biomedical graphs from heterogeneous sources. The KG incorporated entities from multiple OBO ontologies, including:

- Diseases from Mondo Disease Ontology
- Phenotypes from the Human Phenotype Ontology
- Anatomical structures from Uberon
- Gene Ontology (biological processes, cellular components, molecular functions)
- Proteins from the Human Protein Ontology
- Chemical compounds from ChEBI
- Genetic features from Sequence Ontology
- Cellular types from Cell Ontology and Cell Line Ontology

The pipeline included three core stages: *Ontology Unification, Data Curation*, and *KG Assembly*. During unification, potential discrepancies in ontology syntax (e.g., duplicate entries, deprecated terms) were resolved. Data curation involved mapping identifiers and preparing edge relationships. The final stage generated the node-edge structure and assembled the semantically enriched KG.

Additionally, the original PheKnowLator pipeline was extended to encompass:

- Ontology of Adverse Events
- Drug Interaction Knowledge Base (DIKB): supporting evidence for enzyme interactions, including results from clinical and laboratory studies
- DrugCentral: documented drug-transporter interactions, bacterial interactions, and enzyme modulations

FDA Drug Interaction Database: pharmacokinetic outcomes from in vitro and human experiments, retaining only interactions with a fold-change ≥ 2.0 in area under the curve (AUC)

This twofold threshold, mathematically represented as:

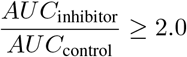

was used to denote interactions with a measurable pharmacokinetic impact.

The above data sources were incorporated into the KG using the OWL-NETS method, which transforms Web Ontology Language (OWL) constructs into biologically meaningful edges. Furthermore, a time-stratified approach was adopted to facilitate longitudinal analysis, enabling predictions about future interactions based on historical data.

### B. Evaluation

A unified heterogeneous graph, named NP-KG, was developed by integrating the ontology-derived knowledge graph with the literature-mined graph using the NetworkX multidigraph framework. This merged graph underwent analytical assessment through focused case studies involving green tea and kratom. Furthermore, an error investigation was conducted to detect discrepancies arising from relation extraction processes and conflicting assertions within NP-KG.

#### 1) Reference Benchmark

Pharmacokinetic interactions confirmed via laboratory experiments or clinical trials were adopted as the reference benchmark for validation. In the absence of clinical trial evidence, outcomes derived from in vitro evaluations and mechanistic simulations were deemed sufficient. If conflicting data emerged between in vitro and clinical findings, the latter were prioritized. The benchmark dataset was curated from an expert-validated pharmacological repository containing experimental and clinical outcomes regarding natural product-drug interactions.

#### 2) Assessment Techniques

## I. Knowledge Reacquisition

To determine the extent to which NP-KG reconstructs established relationships between green tea or kratom and specific metabolic enzymes and transport proteins, identifiers for enzymes and transporters from an ontology-compliant database were initially retrieved. Subsequently, for entities corresponding to green tea (including constituents such as epicatechin3-gallate, epigallocatechin gallate, epicatechin, catechin, and gallocatechin) and kratom (including the alkaloid mitragynine), direct edges and minimal-length connection paths to the metabolic entities were extracted from NP-KG.

The derived connections were systematically compared against the established benchmark data, considering effects such as inhibition, induction, and lack of interaction. Agreement, contradiction, and omission were computed as follows:

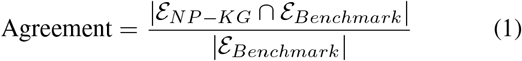

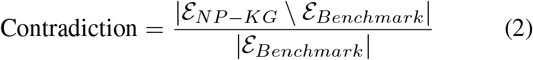

A match was defined when the link or shortest path in NP-KG mirrored the reference outcome. A contradiction arose if NP-KG’s edge opposed the benchmark evidence. Mixed cases were manually examined using metadata embedded in NP-KG. Relations for which congruity or contradiction could not be concluded, such as `interacts_with`, `molecularly_interacts_with`, and `directly_regulates_activity_of`, were excluded from the evaluation metrics. Bidirectional shortest path algorithms provided the path calculations using NetworkX.

### II. Meta-path Inference

Meta-paths, sequences of semantically associated nodes and edges, are utilized for hypothesis development within structured biological graphs. In this context, NP-KG was examined for potential pharmacokinetic interaction pathways involving green tea and kratom using both direct and meta-path traversal techniques. As illustrated in Fig. 4, candidate pathways were obtained by retrieving neighborhood subgraphs surrounding the natural product and constituent nodes and filtering for nodes and relations pertinent to drugs, enzymes, and transporters.

**Fig. 1.**
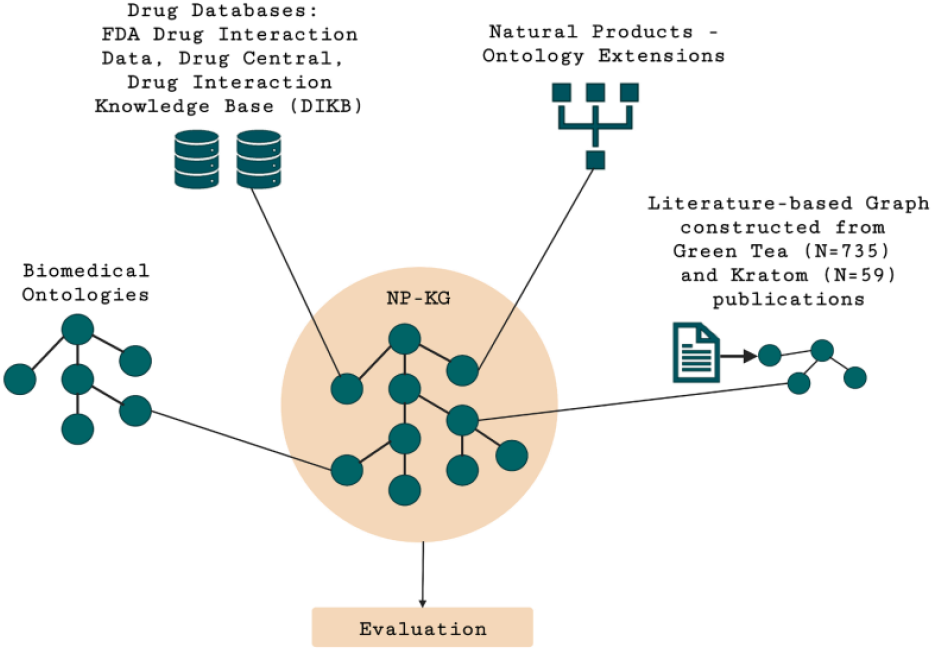
Schematic representation of NP-KG integrating biomedical ontologies, regulatory drug databases, natural products, and literature-derived connections related to green tea and kratom.

**Fig. 2.**
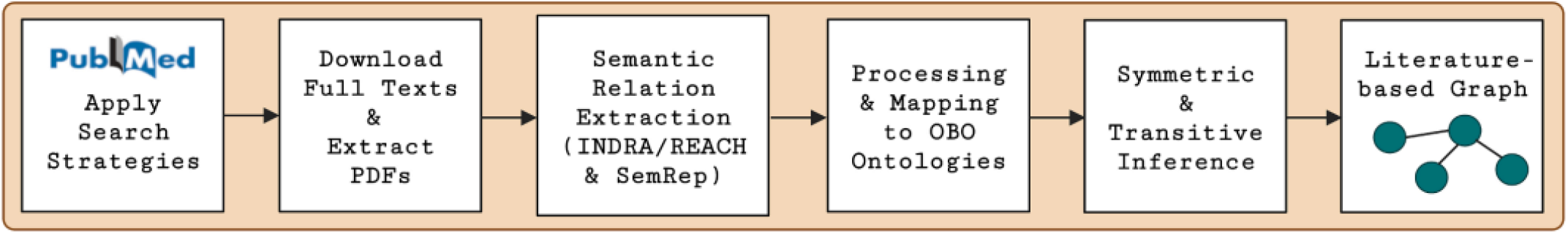
Workflow for semantic relation extraction and literature-based graph construction.

**Fig. 3.**
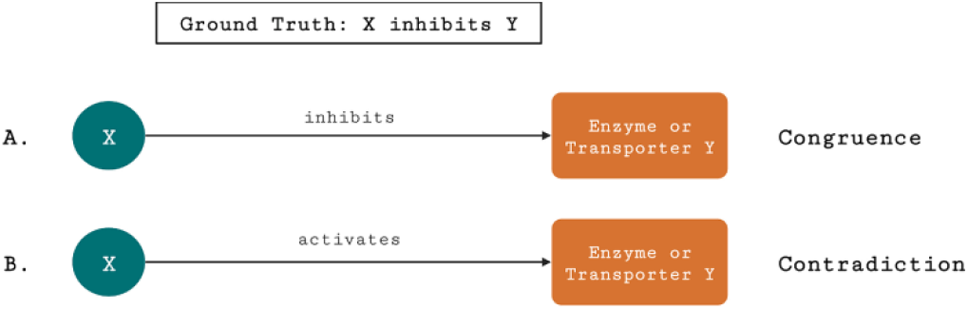
Illustration of agreement and contradiction scenarios for node *X* interacting with enzyme or transporter *Y* in NP-KG, where the reference indicates inhibition.

**Fig. 4.**
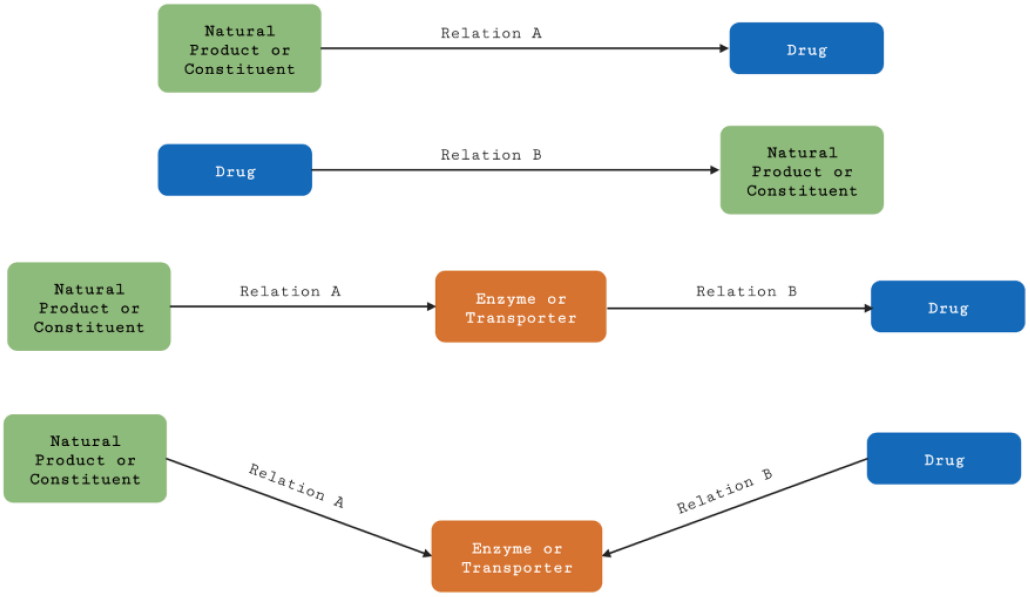
Meta-path and direct edge extraction strategy applied to NP-KG for detecting pharmacokinetic interactions between natural products, enzymes, and drugs.

The extracted subgraphs adhered to specified relational criteria, including but not limited to: `interacts_with`, `molecularly_interacts_with`, `associated_with`, `directly_regulates_activity_ inhibits`, `regulates_activity_of`, and `transports`. These selected paths were subjected to qualitative validation against benchmark information.

Five interaction pairs were prioritized for meta-path analysis due to existing experimental or anecdotal reports: green tea–raloxifene, green tea–nadolol, kratom–midazolam, kratom–quetiapine, and kratom–venlafaxine. These pairs were selected based on documented effects such as decreased systemic exposure, altered intestinal absorption, or cytochrome P450 inhibition by kratom alkaloids. Case-specific evidence for each was traced through the graph to uncover plausible interaction mechanisms.

### III. Knowledge Regeneration

Table I outlines the harmonized and conflicting associations derived from NP-KG in relation to pharmacokinetic interactions involving botanical agents and biochemical mediators, such as enzymes and transport proteins, when aligned with verified reference datasets. Regarding green tea-associated vertices, a total of 59 queries were conducted to identify either direct connections or the shortest relational trajectories across NP-KG that incorporated 19 enzymes and 8 transport elements. Correspondingly, 14 queries were initiated for kratom-linked vertices, encompassing 10 enzymes and a single transporter. Examples of these connectivity patterns, including direct relationships and minimal paths, are visualized in Fig. 5. Numerous vertices revealed multiple relational connections, such as those observed between catechin and UGT1A1 (Fig. 5a), mitragynine and CYP3A (Fig. 5d), and mitragynine and CYP2D6 (Fig. 5e). Both concordant and divergent linkages among these vertices were subjected to manual assessment for validation of either alignment or conflict. Manual evaluations produced the following insights:

**Fig. 5.**
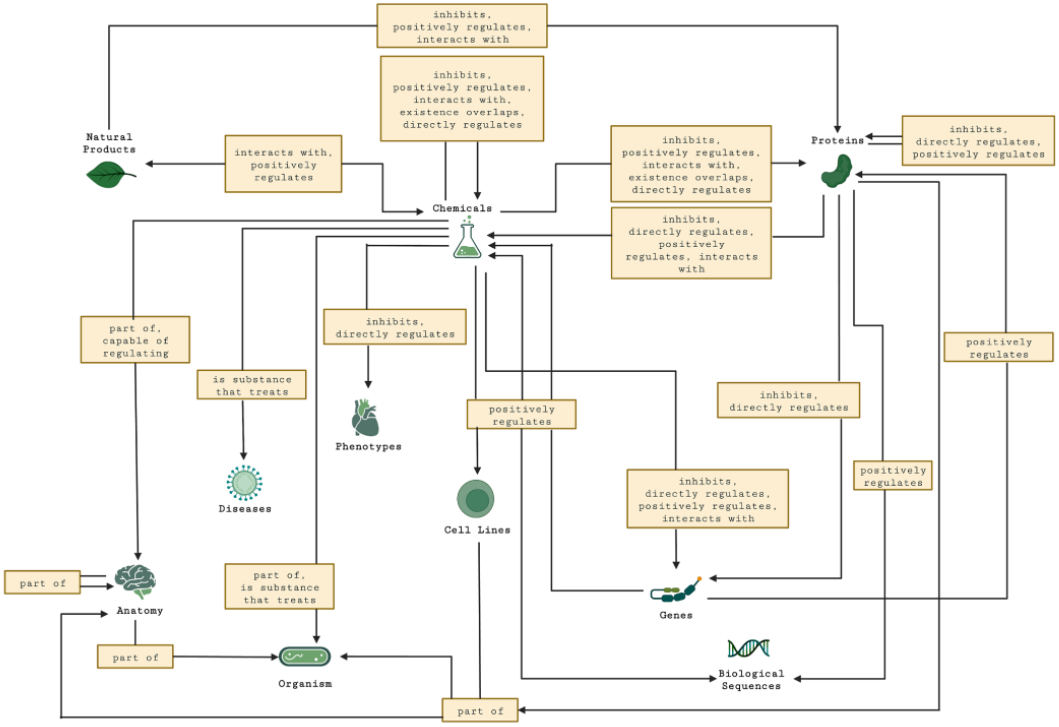
Sample direct and shortest-path connections within NP-KG involving enzymes and natural products.

**TABLE I.** Summary of Interaction Validation from Knowledge Recapturing.

| Botanical | Total Queries | Unresolved (%) |
| --- | --- | --- |
| Green Tea | 59 | 25 (42.37%) |
| Kratom | 14 | 3 (21.43%) |

- The interactions annotated as *inhibits, positively regulates*, and *directly positively regulates quantity of* between EGCG and CYP1A2, as well as EGCG and UGT1A1, demonstrated indications of both consistency and disparity. Contradictions were attributable to divergent conclusions presented in scientific texts.
- Mitragynine, the primary indole-based alkaloid in kratom, functions as a time-dependent suppressor of CYP3A and a reversible antagonist of CYP2D6. The NP-KG paths incorporating *inhibits* and *directly positively regulates quantity of* between mitragynine and CYP3A4 (Fig. 5d) suggested both affirmation and refutation in comparison to the control data. Inspection of source statements revealed reports indicating both inhibition and induction of CYP3A4 in laboratory experiments.
- The presence of both inhibitory and activating associations between mitragynine and CYP2D6 (Fig. 5e) implied incongruent alignment with the baseline. This contradiction was traced to inaccuracies in relational extraction, particularly for the *positively regulates* association.

In approximately 42.37% of searches related to green tea nodes (25 instances) and 21.43% of kratom node queries (3 instances), it was infeasible to determine alignment or contradiction due to the presence of generalized interaction labels such as *interacts with* or *regulates activity of*. For example, NP-KG recorded an *interacts with* association between mitragynine and CYP2C19, although empirical data indicated inhibition. Consequently, these results lacked conclusiveness. Supplementary File 1 contains all paths and associations evaluated for this knowledge regeneration process for both green tea and kratom entities.

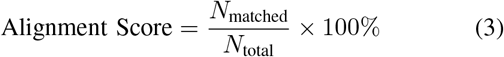

## IV. Meta-path Identification

Through querying of direct associations and intermediate relational chains, the NP-KG was utilized to uncover potential pharmacokinetic natural product–drug interactions (NPDIs) involving green tea and kratom, as presented in Fig. 6. Table II encapsulates findings from both direct and meta-path explorations involving the following pairs: green tea–raloxifene, green tea–nadolol, kratom–midazolam, kratom–quetiapine, and kratom–venlafaxine.

**Fig. 6.**
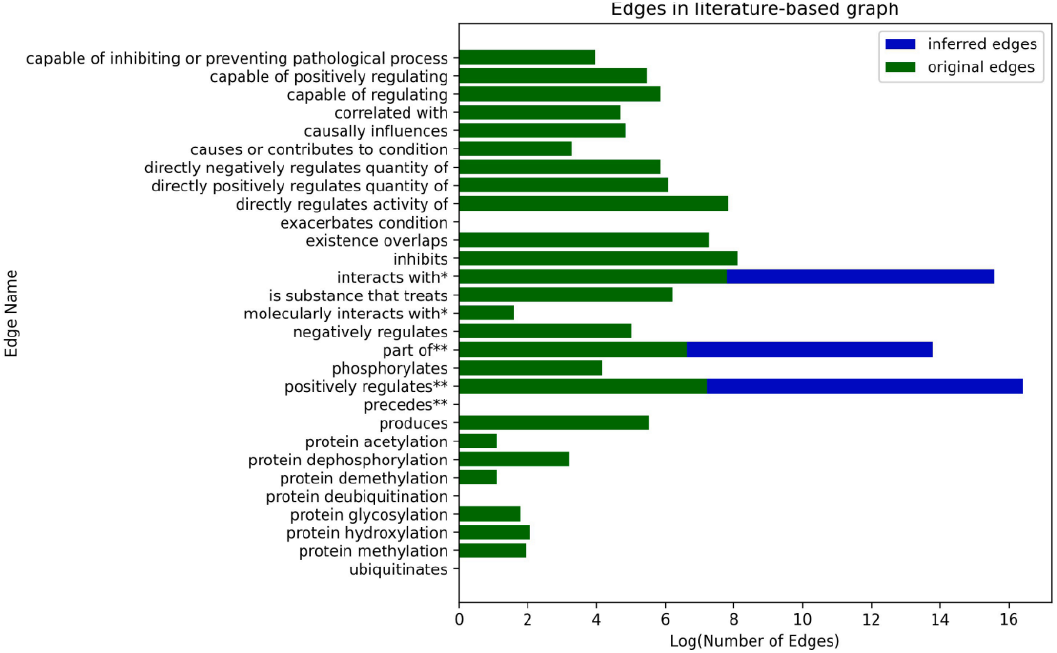
Pharmacokinetic interaction discovery through direct and meta-path queries in NP-KG.

**TABLE II.** Meta-path Discovery Results Across NP-KG.

| NP-Drug Pair | Enzymes Identified | Transporters Identified |
| --- | --- | --- |
| Green Tea-Raloxifene | CYP3A4, UGTs | None |
| Green Tea-Nadolol | 8 Enzymes | 12 Transporters |
| Kratom-Midazolam | CYP3A4, CYP2D6 | P-gp |
| Kratom-Quetiapine | CYP3A | None |
| Kratom-Venlafaxine | CYP3A, CYP2D6 | None |

**TABLE III.** Overview of alignment and divergence in NP-KG against validated sources.

|  | Green Tea (%) | Kratom (%) |
| --- | --- | --- |
| Concordance | 23 (38.98) | 7 (50.0) |
| Discrepancy | 9 (15.25) | 3 (21.43) |
| Only paths/edges found | 25 (42.37) | 3 (21.43) |
| Both cases observed | 2 (3.39) | 1 (7.14) |

**TABLE 4.** Meta-path analysis summary for select natural product-drug pairs. (ND = Not Detected)

| Pair | Enzyme(s) | Transporter(s) |
| --- | --- | --- |
| Green tea-raloxifene | CYP3A4, UGT | ND |
| Green tea-nadolol | See Appendix A.4 | See Appendix A.5 |
| Kratom-midazolam | CYP2D6, CYP3A4 | P-glycoprotein |
| Kratom-quetiapine | CYP3A4 | P-glycoprotein |
| Kratom-venlafaxine | CYP2D6, CYP3A4 | ND |

### A. Green Tea–Raloxifene Interaction

Raloxifene, an anti-osteoporotic agent and substrate of intestinal UGTs, displayed concentration-dependent suppression of UGT function in vitro due to exposure to green tea constituents. However, in a subsequent human study, a 30% decline in the area under the plasma concentration–time curve (AUC) was recorded when comparing green tea to water. This reduction opposed initial in vitro predictions, which anticipated an AUC increase. No substantial variation was detected in raloxifene’s half-life or glucuronide-to-parent drug AUC ratios, thereby excluding intestinal UGT inhibition as the mechanism of interaction. NP-KG uncovered links involving UGTs and CYP3A4 for this interaction, while CYP2C9 was connected to raloxifene but not to green tea. Fig. 7 offers a visual representation of these associations.

**Fig. 7.**
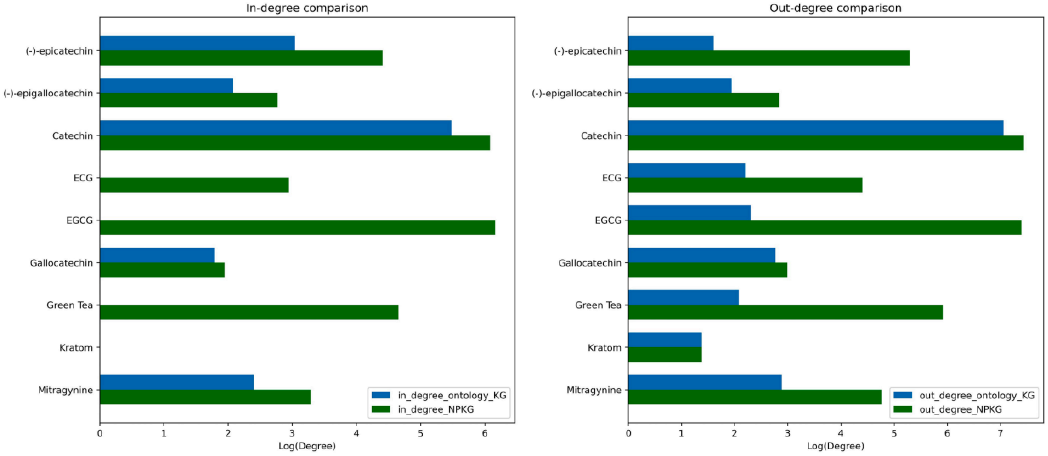
Interaction network between green tea components and raloxifene in NP-KG.

**Fig. 8.**
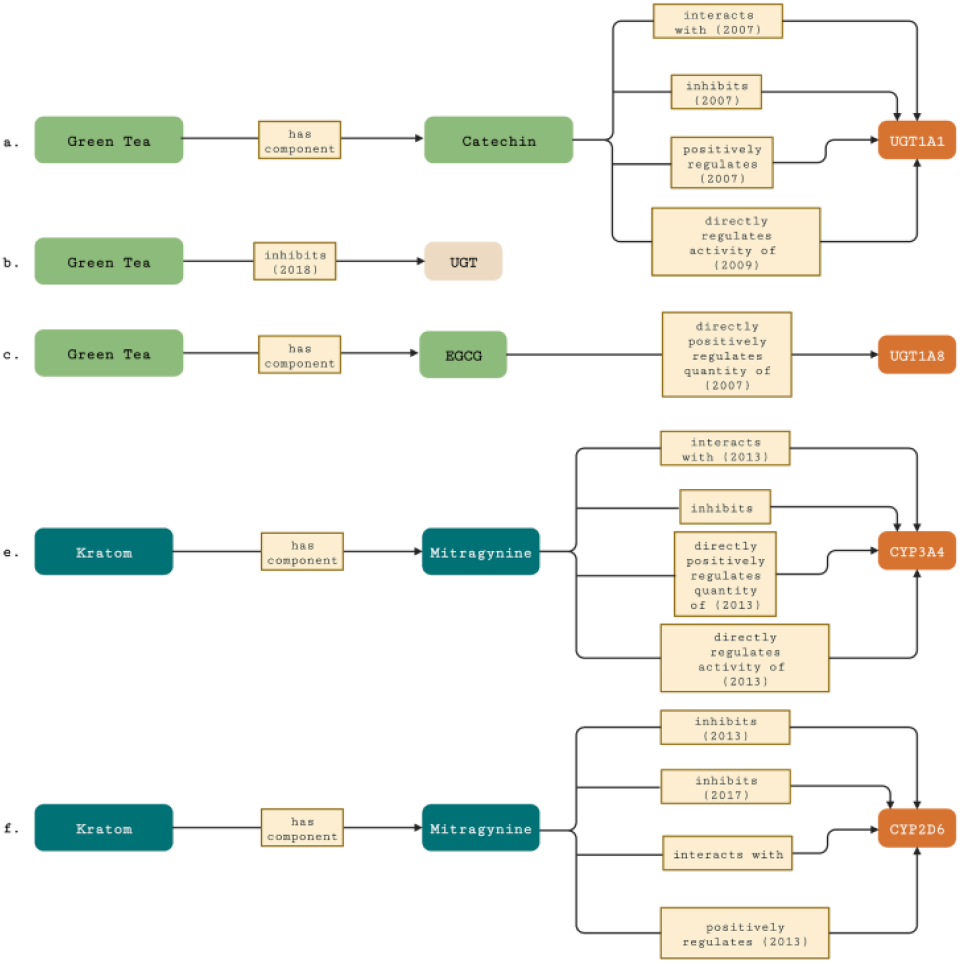
Node degree comparison for green tea and kratom between ontologyderived KG and NP-KG. (a) In-degree distribution; (b) Out-degree distribution. Horizontal axis shows logarithmic degree values, vertical axis displays node labels.

**Fig. 9.**
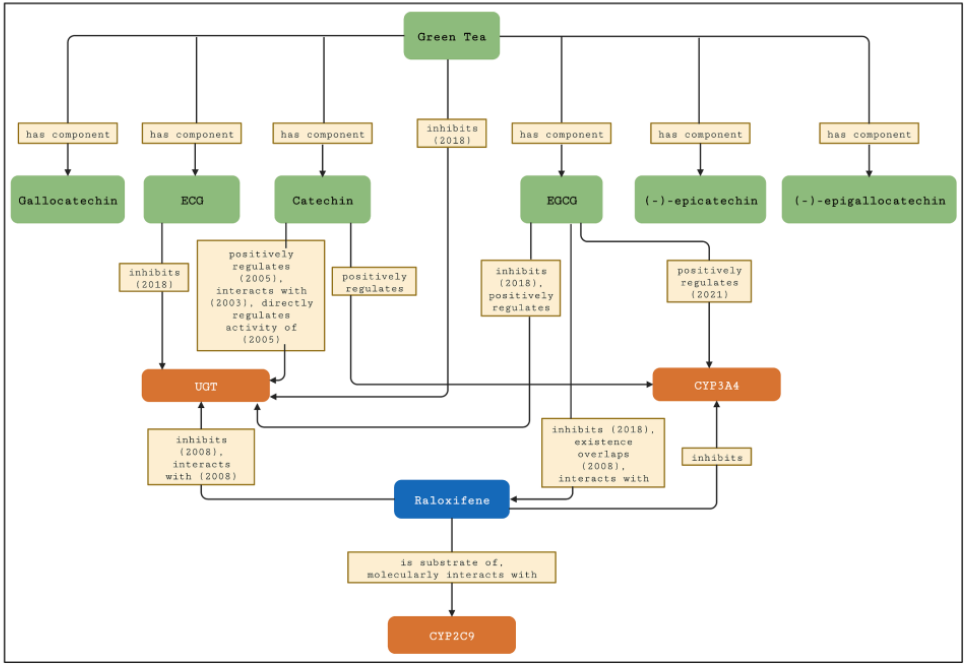
NP-KG subgraph of pharmacokinetic pathways involving green tea and raloxifene with CYP3A4, CYP2C9, and UGT. Nodes and edges are annotated with literature-derived metadata including publication year.

**Fig. 10.**
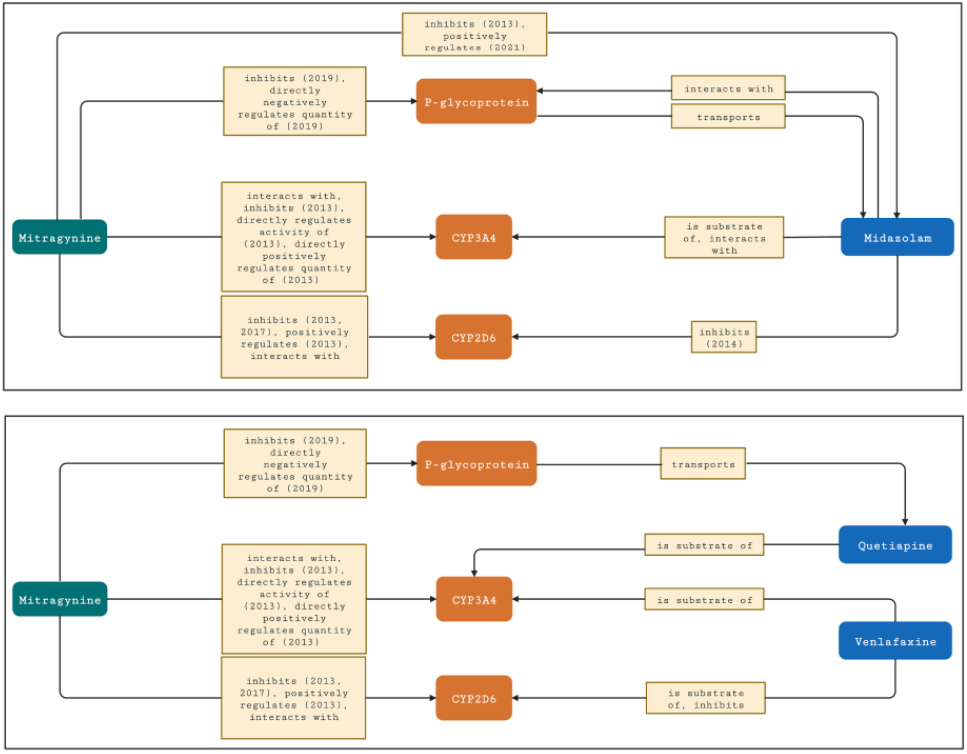
NP-KG subnetwork for mitragynine-midazolam interaction highlighting CYP3A4 and CYP2D6 with annotated literature-derived edges.

### B. Green Tea–Nadolol Interaction

Nadolol, a non-selective *β*-adrenergic blocker employed in hypertension management, demonstrated a significant pharmacokinetic alteration when co-administered with green tea. The beverage reduced nadolol’s AUC by 80% and diminished its therapeutic blood pressure-lowering impact. Although laboratory tests showed nadolol is transported by OATP1A2 and that EGCG suppressed OATP1A2 uptake, the role of this transporter remains debated. Other mechanisms, such as increased efflux transporter activity, may be involved. NPKG meta-paths highlighted 8 enzymes and 12 transporters as contributors to this interaction, detailed in Appendix Tables A.4 and A.5.

### C. Kratom–Midazolam Interaction

Midazolam, a benzodiazepine metabolized primarily via CYP3A, is susceptible to interaction with kratom constituents. Mitragynine and kratom extracts were found to inhibit CYP2C9, CYP2D6, and CYP3A isoforms in cell-based models. A mechanistic model predicted a pharmacokinetic disruption due to time-sensitive CYP3A inhibition by mitragynine. Meta-path analysis through NP-KG identified two enzymes—CYP3A4 and CYP2D6—and the transporter P-glycoprotein as mediators of this interaction. Nevertheless, midazolam is not considered a substrate for either CYP2D6 or P-glycoprotein, suggesting indirect mechanisms.

### D. Kratom–Quetiapine and Kratom–Venlafaxine Interactions

Venlafaxine, an antidepressant acting through CYP3A and CYP2D6 metabolism, and quetiapine, an antipsychotic processed via CYP3A, have both been reported to exhibit interaction risk when consumed alongside kratom. Proposed mechanisms involve enzyme inhibition by mitragynine. Metapath analysis confirmed the involvement of both CYP3A and CYP2D6 within these pharmacokinetic interplays, substantiating the predicted interactions.

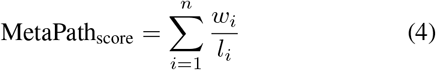

## V. Discussion

This research introduces a unique integration of structured biomedical ontologies and machine-extracted insights from comprehensive scientific texts to develop a knowledge graph (KG) centered on natural product-drug interactions (NPDIs). The resulting NP-KG is distinct in its fusion of domain ontologies, accessible biomedical repositories, and specialized textual information to propose mechanistic interpretations for pharmacokinetic NPDIs. The framework enables the reinterpretation of existing pharmacokinetic relationships while offering a framework for uncovering lesser-known interactions, thus facilitating the strategic formulation of new experimental designs.

Two assessment methodologies were employed to verify the inferred NPDI mechanisms involving green tea and kratom. The initial strategy involved identifying immediate connections and shortest relational sequences between natural product nodes and associated metabolic enzymes or transport proteins within the NP-KG. A juxtaposition with the NaPDI Center’s reference data revealed both consistencies and discrepancies. Follow-up scrutiny attributed these discrepancies to inconsistencies in the source literature and inaccuracies in automated relation extraction processes. In several instances, both agreement and disagreement appeared for the same query, necessitating manual inspection (15.25% for green tea, 21.43% for kratom).

Analyzing contradictions in literature-derived biomedical KGs is an evolving domain. Prior investigations into semantic inconsistencies in databases like SemMedDB have uncovered multiple causative elements including variations in study cohorts, scientific debates, evolving knowledge, and methodological differences. Detection of such discrepancies frequently necessitates examination of not only the sentence-level but also broader abstract-level context.

The second validation technique applied meta-path queries and direct edges to specific product-drug combinations: green tea-raloxifene, green tea-nadolol, kratom-midazolam, kratom-quetiapine, and kratom-venlafaxine. Meta-paths, which represent multi-hop connection patterns, offer a way to hypothesize biological mechanisms underlying interactions. Analysis indicated associations such as UGT and CYP3A4 in green tea-raloxifene interaction, although clinical data did not always support the inferred mechanisms. This illustrates the potential gap between in vitro observations and in vivo outcomes.

The identification of OATP1A2 in the green tea-nadolol pathway points to the necessity of exploring additional pathways. NP-KG inferred the involvement of major enzymes and transporters in kratom-drug combinations, several of which align with existing research, whereas others remain to be substantiated.

The construction of the KG was complicated by limited access to machine-readable full texts and heterogeneity in terminology across ontologies, databases, and extraction tools. While prior drug-related KGs often relied exclusively on curated datasets or text mining, this approach incorporated both, utilizing over 700 full-text scientific documents with outputs from two independent relation extraction pipelines.

The degree analysis between ontology-based and literature-enhanced KGs indicated that NP-KG offers more detailed connectivity for green tea and kratom. The inclusion of textual predications filled gaps in ontology-only graphs. A significant volume of data related to green tea reflects its widespread use and higher representation in published literature compared to kratom.

Integration of semantic triples using instance-level representation facilitated accurate inference. The harmonization of extracted relationships into OBO-compliant terms further streamlined integration. While some inaccuracies in extraction remain, improvement in these tools could enhance graph accuracy.

The overarching objective remains to identify overlooked pharmacokinetic NPDIs and generate plausible mechanistic insights with corroborative evidence. This can be achieved by expanding NP-KG with additional products and utilizing:

- Meta-path based symbolic reasoning;
- Hypothesis generation using spontaneous reporting system data;
- Graph representation learning methods for KG completion and link prediction.

These approaches collectively hold promise in bridging knowledge deficits and uncovering complex pharmacological interactions involving underexplored natural compounds. Quantitatively, the meta-path analysis revealed that 4 out of 5 examined NP–drug pairs included at least one interaction not catalogued in benchmark datasets, suggesting the framework’s potential to generate novel, testable hypotheses with an estimated discovery yield of 80% in the current evaluation.

## VI. Limitations and Challenges

This study presents several inherent constraints. Primarily, the literature retrieval procedures did not involve manual refinement, potentially introducing irrelevant semantic assertions into the text-derived knowledge graph. Additionally, the literature-based graph encompasses known inaccuracies stemming from suboptimal entity recognition, omissions due to low recall, and challenges in parsing intricate sentence structures. Prior assessments have estimated the recall rate of abstract-based relation extraction using SemRep to be approximately 42%, though internal evaluations indicated even lower recall levels for pharmacokinetic full texts when benchmarked against manually curated datasets. Notably, SemRep has not undergone formal validation for full-text documents.

Another limitation arises from the non-targeted extraction of predications—relation extraction was performed on entire texts without isolating specific structural components such as tables or methodologically critical sections. Moreover, the current mechanism for linking entities within literature-derived predications achieved only 60–70% mapping accuracy between identified subjects and objects, which may have resulted in loss of pertinent relational data in NP-KG.

Improvements could involve integrating domain-adapted named entity recognition and relation extraction tools—customized for natural products—along with supplementary knowledge bases tailored to dietary supplements and food-derived compounds. These enhancements would augment the coverage of underrepresented entities and strengthen linkage fidelity.

Furthermore, while meta-path and shortest-path algorithms were employed for knowledge inference and graph evaluation due to their computational simplicity, their scope remains constrained. These symbolic techniques, for instance, did not account for downstream metabolites or intermediate pharmacokinetic processes that may elucidate the biochemical mechanisms behind specific NPDIs.

Embedding-based methodologies, such as those derived from graph representation learning, could address these limitations by generating vectorized representations of KG nodes and edges. This would enable the extrapolation of new, unobserved connections within the graph space and extend inference capabilities beyond symbolic pathways.

Finally, establishing the validity of inferred NPDI mechanisms presents a significant challenge, given the limited availability of verified benchmark data. Unlike drug–drug interactions, which are extensively catalogued, few structured resources contain mechanistic data for natural product–drug interactions. Existing databases related to supplement interactions provide only partial validation support. As such, much of the evaluation must rely on manually curated literature evidence or indirect adaptation of drug–drug interaction frameworks for food–drug interaction scenarios.

In this investigation, validation efforts centered on green tea and kratom, utilizing enzyme and transporter associations from the NaPDI Center database. The scope of evaluation was confined to interactions supported by existing reference data. However, NP-KG also possesses the capacity to assist in discovering beneficial natural product–drug interactions not covered by this analysis. Future investigations could expand target scope to include nuclear receptors and other pharmacokinetic mediators, offering deeper insights into these multifaceted interactions. To overcome these shortcomings, future iterations of NP-KG will incorporate domain-adapted named entity recognition and relation extraction tools, such as BioBERT-based models or tools tailored to pharmacokinetic literature. These alternatives demonstrate improved recall and precision in extracting drug-enzyme-transporter interactions, especially from full-text articles. In addition to symbolic inference methods, we plan to integrate graph representation learning techniques such as TransE, RotatE, and Graph Convolutional Networks (GCNs) to derive latent relationships within NP-KG. These embedding models can enhance prediction accuracy by capturing global graph structure and semantic proximity, offering a powerful complement to explicit meta-path discovery. Incorporating structured document elements—such as tables, figure captions, and methods sections—during relation extraction can enhance the quality of extracted knowledge by capturing more specific and reliable assertions that are often missed by sentence-based parsing alone. Observed relation extraction errors were frequently attributed to complex biomedical sentence constructions, the presence of negation cues, coreference ambiguity, and use of semantically overloaded terms. Enhanced parsing strategies and biomedical-specific language models could help mitigate these challenges.

## VII. Conclusions

This work introduced NP-KG, a knowledge graph that synthesizes curated biomedical ontologies and databases with semantically enriched full-text scientific documents to investigate natural product–drug interactions (NPDIs). By integrating structured vocabularies with machine-extracted predications, NP-KG offers a hybrid infrastructure capable of generating hypotheses supported by both established pharmacological knowledge and new evidence drawn from the biomedical literature.

The utility of NP-KG was demonstrated through targeted evaluations involving green tea and kratom, revealing its capability to propose plausible pharmacokinetic mechanisms for interactions involving enzymes and transporters. These insights underscore NP-KG’s potential to streamline hypothesis prioritization in preclinical research focused on identifying clinically meaningful NPDIs.

Looking forward, enhancements are planned to improve relation extraction performance, broaden the representation of natural products in the KG, and incorporate advanced inference strategies—such as graph embedding models—for robust hypothesis generation. Expansion efforts include the addition of novel natural products and integration of auxiliary biomedical knowledge domains to further increase the graph’s utility. To further generalize NP-KG’s applicability, future work will expand its scope to include additional commonly used natural products such as St. John’s Wort, turmeric, ginseng, and piperine. These additions will allow more comprehensive evaluation and refinement of the framework across diverse NPDI scenarios. To support wider adoption by clinical researchers and pharmacologists, the framework will also include interactive visualization dashboards. These will enable users to explore meta-paths, trace evidence sources, and contribute feedback, thereby facilitating iterative refinement of the graph. While this study focused on identifying potential conflicts, the same framework can be adapted to uncover beneficial natural product–drug synergies, such as bioavailability enhancement through transporter modulation or enzyme inhibition.

